# An 8-Gene Bevacizumab Resistance Signature Predicts Prognosis and Reveals Immunosuppressive Microenvironment in Colorectal Cancer

**DOI:** 10.64898/2026.05.17.725749

**Authors:** Niu Zhengchuan, Qiu Dongze, Xu Pingping

## Abstract

**Background:** Bevacizumab resistance severely limits long-term efficacy in metastatic colorectal cancer (CRC). This study aimed to develop and validate a bevacizumab resistance-associated gene signature for prognosis prediction and immune microenvironment characterization in CRC.

**Methods:** Two GEO datasets (GSE19862, GSE86582) with bevacizumab response data and TCGA-COAD/READ RNA-seq data were analyzed. Overlapping differentially expressed genes (DEGs) linked to both CRC progression and bevacizumab resistance were identified. An 8-gene signature (AXIN2, PSORS1C1, KRT74, SLC2A3, STIL, IL33, GALNT6, HSD11B2) was constructed via univariate Cox and LASSO-Cox regression.

**Results:** In the TCGA cohort, high-risk patients had shorter overall survival (OS; log-rank *P* < 0.0001). Time-dependent ROC yielded 1-year AUC = 0.638, 3-year AUC = 0.657, and 5-year AUC = 0.757. Multivariate Cox regression confirmed the risk score as an independent prognostic factor. External validation in GSE39582 (optimal cutoff = −1.49) replicated these findings: high-risk patients had inferior OS (*P* = 0.0016) with acceptable 1/3/5-year AUCs and retained independent prognostic value (HR = 1.634, *P* = 0.00415). CIBERSORT and ESTIMATE analyses showed that the high-risk group was characterized by increased M2 macrophages and neutrophils, higher immune and stromal scores, and reduced activated memory CD4^+^ T cells, monocytes, and activated dendritic cells (all *P* < 0.05). GSEA highlighted enrichment of TNF-α/NF-κB, IL-6/JAK/STAT3, and immune checkpoint pathways in the high-risk group. AXIN2 (HR = 0.829, *P* = 0.032) was an independent protective factor, while PSORS1C1 (HR = 1.356, *P* = 0.048) was an independent risk factor.

**Conclusion:** The 8-gene bevacizumab resistance signature robustly predicts prognosis and reflects an immunosuppressive microenvironment closely linked to bevacizumab failure in CRC. These findings provide novel insights into immune-mediated resistance and support clinical risk stratification.

## 1 Introduction

Colorectal cancer (CRC) is one of the most prevalent malignancies worldwide and a leading cause of cancer-related death [1]. For metastatic CRC, bevacizumab combined with chemotherapy is standard first-line therapy [2]. However, intrinsic and acquired resistance markedly reduces long-term efficacy, and reliable predictive biomarkers remain lacking.

The tumor immune microenvironment (TIME) critically influences therapeutic response and prognosis. Accumulating evidence indicates that an immunosuppressive microenvironment, chronic inflammation, and stromal remodeling are tightly associated with bevacizumab resistance [3,4]. Nonetheless, the underlying mechanisms remain incompletely understood.

Transcriptome-based bioinformatics has been widely applied to identify prognostic signatures and explore molecular mechanisms. Although numerous prognostic models have been developed for CRC [5–7], few are directly derived from bevacizumab resistance profiles and integrated with immune landscape characterization.

Here, overlapping DEGs from two bevacizumab-resistant GEO datasets and the TCGA cohort were screened to construct an 8-gene prognostic signature via LASSO-Cox regression. The model was validated in both TCGA and an independent external cohort. Immune infiltration, ESTIMATE, and GSEA analyses were performed to explore biological functions and immune-related mechanisms. The study workflow is summarized in the flowchart. This work identifies a robust prognostic biomarker and provides novel insights into bevacizumab resistance in CRC by focusing on the immunosuppressive TIME.

## 2 Materials and Methods

### 2.1 Data Collection and Preprocessing

CRC gene expression and clinical data were retrieved from GEO and TCGA databases. Two bevacizumab-related GEO datasets (GSE19862, GSE86582) with treatment response data were used for differential analysis. TCGA-COAD and TCGA-READ were integrated into a unified TCGA-CRC cohort after normalization and batch effect removal. GSE39582 was selected as an independent external validation cohort, which contains comprehensive clinicopathological and survival information for CRC patients [8]. Raw data were preprocessed following standard procedures: non-annotated and multi-mapped probes were removed, duplicate gene expressions were averaged, and samples with incomplete survival or clinical data were excluded.

### 2.2 Identification of Differentially Expressed Genes (DEGs)

Two comparisons were performed: bevacizumab-resistant vs. sensitive groups and CRC tumor vs. adjacent normal tissues. In GEO datasets, DEGs were identified using limma (|log_2_FC| > 0.5, *P* < 0.05). In TCGA-CRC, DEGs were screened via edgeR (|log_2_FC| > 0.5, FDR < 0.05). Overlapping genes between the two DEG sets were defined as candidate genes for model construction.

### 2.3 Construction of Prognostic Signature

Univariate Cox regression was applied to screen prognostic genes (*P* < 0.1) using the survival package. LASSO-Cox regression with 10-fold cross-validation was performed via glmnet, and the optimal λ (lambda.min) was selected to identify key genes and build the risk model. The risk score was calculated as: Risk score = Σ(Expression⍰ × LASSO coefficient⍰). Patients were stratified into high-risk and low-risk groups based on the median risk score in the TCGA cohort.

### 2.4 Model Validation

Kaplan–Meier curves and log-rank tests (survminer) were used to compare OS between risk groups. Time-dependent ROC curves (timeROC) assessed 1-, 3-, and 5-year predictive performance. Univariate and multivariate Cox regression confirmed the independent prognostic value of the risk score. GSE39582 was used for external validation with consistent analytical workflows.

### 2.5 Immune Infiltration Analysis

CIBERSORT quantified 22 immune cell subsets using the LM22 matrix (*P* < 0.05) [9]. The ESTIMATE algorithm calculated immune, stromal, and ESTIMATE scores. Spearman’s correlation was used to explore associations between the risk score and immune parameters.

### 2.6 Gene Set Enrichment Analysis (GSEA)

GSEA [10] was performed between risk groups and AXIN2/PSORS1C1 high/low subgroups (NES > 1, *P* < 0.05, FDR < 0.25). Four bevacizumab resistance-related pathways (drug metabolism, DNA damage repair, cell cycle, immune suppression) and a custom immune checkpoint gene set were analyzed. Signature genes were excluded to avoid circular reasoning. Detailed gene sets are provided in Supplementary S1.

### 2.7 Statistical Analysis

All analyses were conducted in R (v4.4.2) with appropriate packages. Wilcoxon rank-sum, χ^2^, Fisher’s exact, log-rank, and Spearman’s correlation tests were performed as applicable. A two-tailed *P* < 0.05 was considered statistically significant.

## 3 Results

### 3.1 Screening of Bevacizumab Resistance-Related Candidate Genes

To identify genes associated with both bevacizumab resistance and CRC progression, three transcriptomic datasets (TCGA-CRC, GSE19862, GSE86582) were analyzed. TCGA-CRC yielded 10,097 DEGs (8,473 unique), GSE19862 yielded 1,458 DEGs (808 unique), and GSE86582 yielded 2,194 DEGs (1,332 unique). Venn diagram analysis identified 60 overlapping DEGs common to all three cohorts (Figure 1A), representing genes involved in both tumor malignancy and drug resistance. These 60 genes were subjected to univariate Cox regression, identifying 9 genes with prognostic significance (*P* < 0.1) (Figure 1B,C): five protective factors (HR < 1: AXIN2, GALNT6, HSD11B2, IL33, STIL) and four risk factors (HR > 1: NDUFA4L2, PSORS1C1, SLC2A3, KRT74). These nine genes were selected for subsequent model construction.

**Figure 1.**
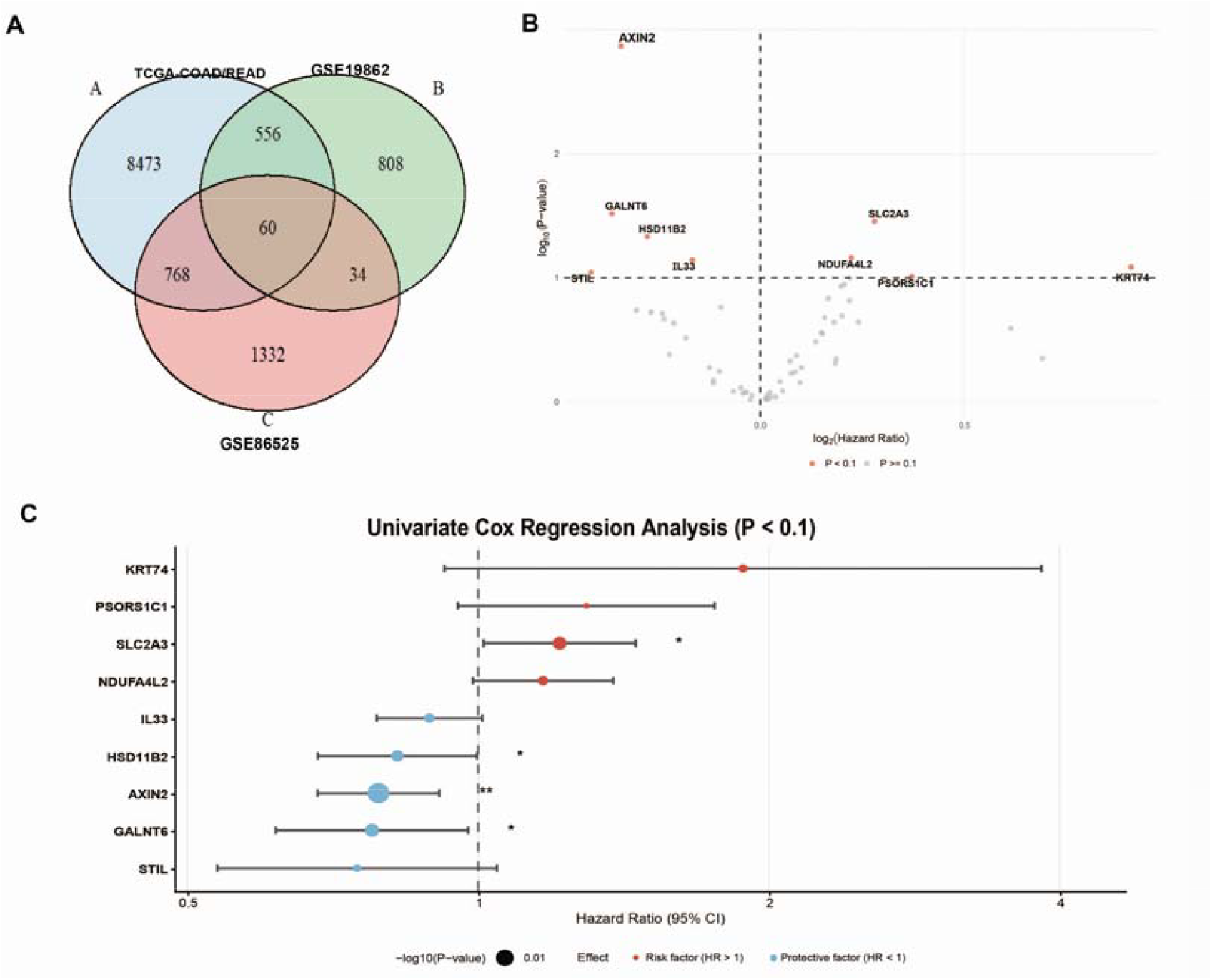
Identification of candidate prognostic genes. (A) Venn diagram showing overlapping DEGs from three datasets (TCGA-COAD/READ, GSE19862, GSE86582). (B) Volcano plot of univariate Cox regression for 60 overlapping genes, with significant genes (*P* < 0.1) highlighted in red. (C) Forest plot of nine genes selected by univariate Cox regression, showing HRs and 95% CIs.

### 3.2 Construction of the 8-Gene Prognostic Signature via LASSO-Cox Regression

To refine the nine candidate genes and build a robust signature, LASSO-Cox regression with 10-fold cross-validation was performed to reduce dimensionality and avoid overfitting. Using the optimal λ (lambda.min) determined by cross-validation (Figure 2A,B), eight genes were retained for model construction, with coefficient trajectories and feature importance shown in Figure 2C: AXIN2 (−0.2305), GALNT6 (−0.1362), HSD11B2 (−0.0741), IL33 (−0.0601), STIL (−0.1879), PSORS1C1 (0.2169), SLC2A3 (0.1685), and KRT74 (0.1386). Multivariate Cox regression confirmed the independent prognostic value of these genes (Figure 2D). The risk score was calculated as a linear combination of gene expression multiplied by corresponding LASSO coefficients.

**Figure 2.**
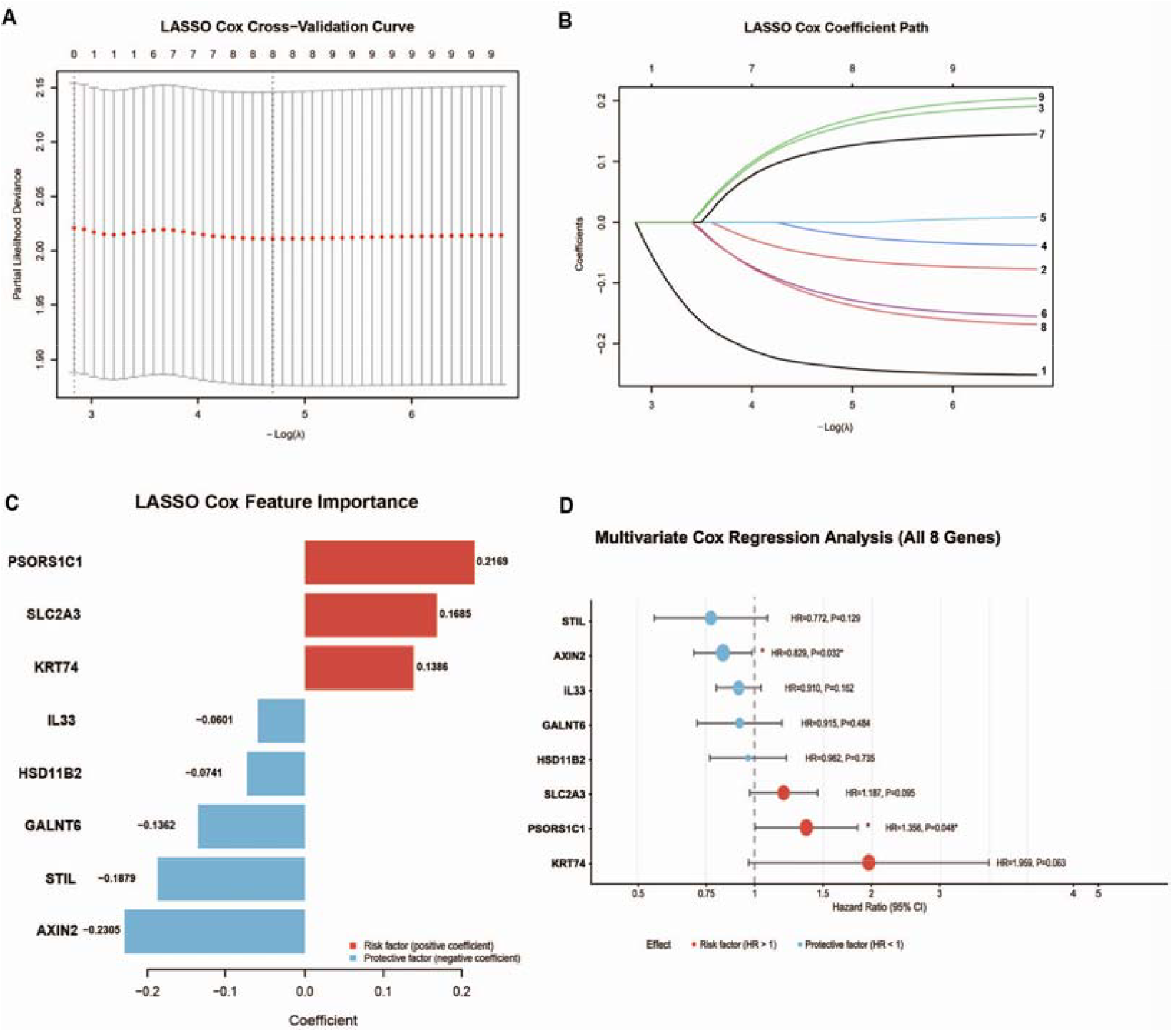
Construction of the 8-gene prognostic signature via LASSO-Cox regression. (A) LASSO Cox cross-validation curve for optimal λ selection. (B) LASSO coefficient profiles of nine candidate genes. (C) Coefficient distribution of eight genes in the final signature. (D) Multivariate Cox regression forest plot for each gene’s independent prognostic value in TCGA.

### 3.3 Prognostic Value of the 8-Gene Signature in the TCGA Cohort

TCGA-CRC patients were stratified into high-risk (n = 238) and low-risk (n = 238) subgroups based on the median risk score (−0.0578) (Figure 3A). Time-dependent ROC curves demonstrated robust predictive performance, with 1-, 3-, and 5-year OS AUCs of 0.638, 0.657, and 0.757, respectively (Figure 3B). Kaplan–Meier analysis revealed significantly shorter OS in the high-risk group (log-rank *P* < 0.0001, Figure 3C). Multivariate Cox regression, adjusted for age, gender, tumor location, and TNM stage, confirmed the risk score as an independent prognostic factor (HR = 2.529, 95% CI: 1.505–4.25, *P* < 0.0005; Figure 3D).

**Figure 3.**
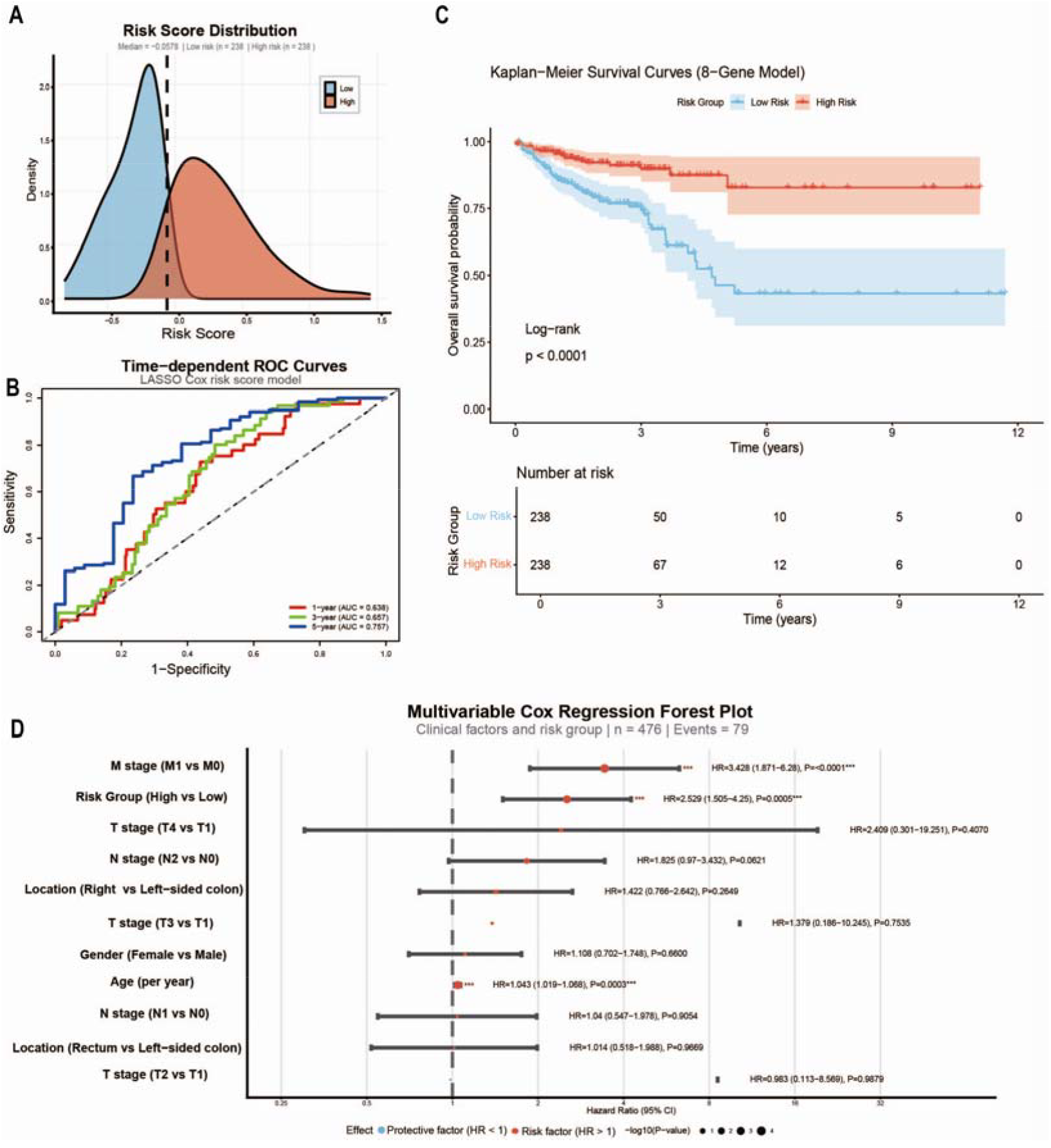
Prognostic performance of the 8-gene signature in the TCGA cohort. (A) Risk score distribution stratified by median cutoff. (B) Time-dependent ROC curves for 1-, 3-, and 5-year survival. (C) Kaplan–Meier curves comparing OS between high- and low-risk groups. (D) Multivariate Cox regression forest plot confirming risk score as an independent prognostic factor.

### 3.4 External Validation of the 8-Gene Signature in the GSE39582 Cohort

To assess robustness and generalizability, external validation was performed in the independent GSE39582 cohort (n = 505). Using the optimal cutoff (−1.49) determined by maximally selected rank statistics (Figure 4A), patients were stratified into high-risk (n = 205) and low-risk (n = 300) subgroups. Time-dependent ROC curves showed good predictive performance, with 1-, 3-, and 5-year AUCs of 0.631, 0.611, and 0.597 (Figure 4B). Kaplan–Meier analysis confirmed inferior OS in the high-risk group (log-rank *P* = 0.0016, Figure 4C), consistent with TCGA findings. Multivariate Cox regression, adjusted for clinical covariates, validated the risk score as an independent prognostic factor (HR = 1.634, 95% CI: 1.177–2.267, *P* = 0.00415; Figure 4D).

**Figure 4.**
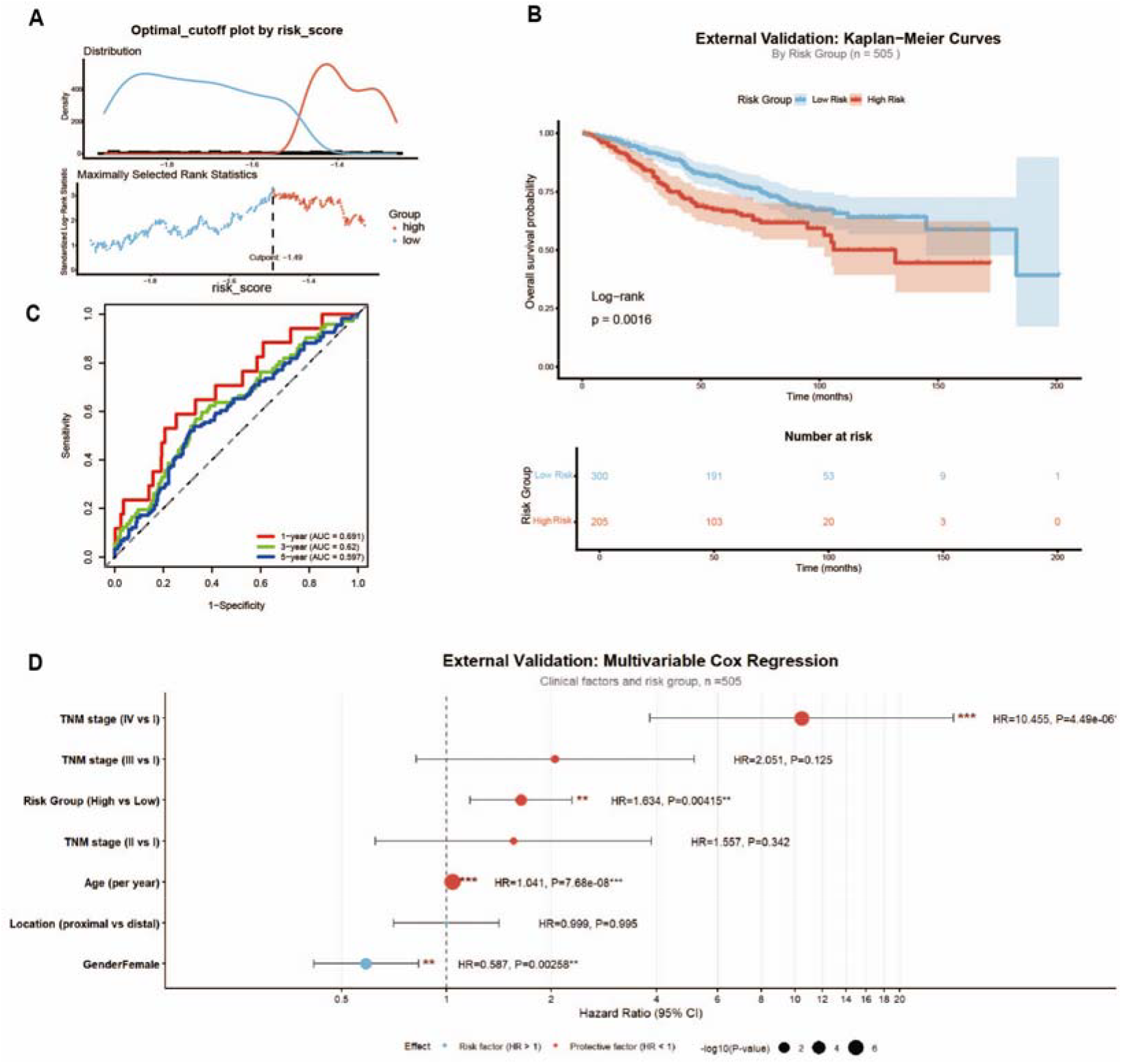
External validation of the signature in the GSE39582 cohort. (A) Optimal risk score cutoff via maximally selected rank statistics. (B) Kaplan–Meier curves for high- and low-risk patients. (C) Time-dependent ROC curves for 1-, 3-, and 5-year survival. (D) Multivariate Cox regression verifying risk score as an independent prognostic factor.

### 3.5 Immune Infiltration Analysis Based on CIBERSORT

To explore the immune landscape associated with the signature, CIBERSORT quantified 22 immune cell subsets in the TCGA cohort. Correlation analysis (Figure 5A) generated a pairwise correlation matrix of immune cell subsets, with color intensity reflecting correlation strength and direction (red: positive; blue: negative). The matrix revealed distinct immune cell co-regulation patterns in TCGA-CRC, underpinning compositional differences between risk groups (Figure 5B). The high-risk group had significantly higher proportions of CD8^+^ T cells, M0/M1/M2 macrophages, and neutrophils (all *P* < 0.05), while resting memory CD4^+^ T cells, plasma cells, activated memory CD4^+^ T cells, monocytes, and activated dendritic cells were markedly reduced (all *P* < 0.05). The high-risk subgroup was characterized by global myeloid expansion and impaired antigen presentation and helper T-cell immunity. Despite higher CD8^+^ T-cell abundance, reduced activated dendritic cells and helper T cells indicated functionally compromised cytotoxic immunity. Collectively, high-risk CRC exhibited a pro-tumorigenic immune landscape with enhanced myeloid inflammation and suppressed adaptive immunity.

**Figure 5.**
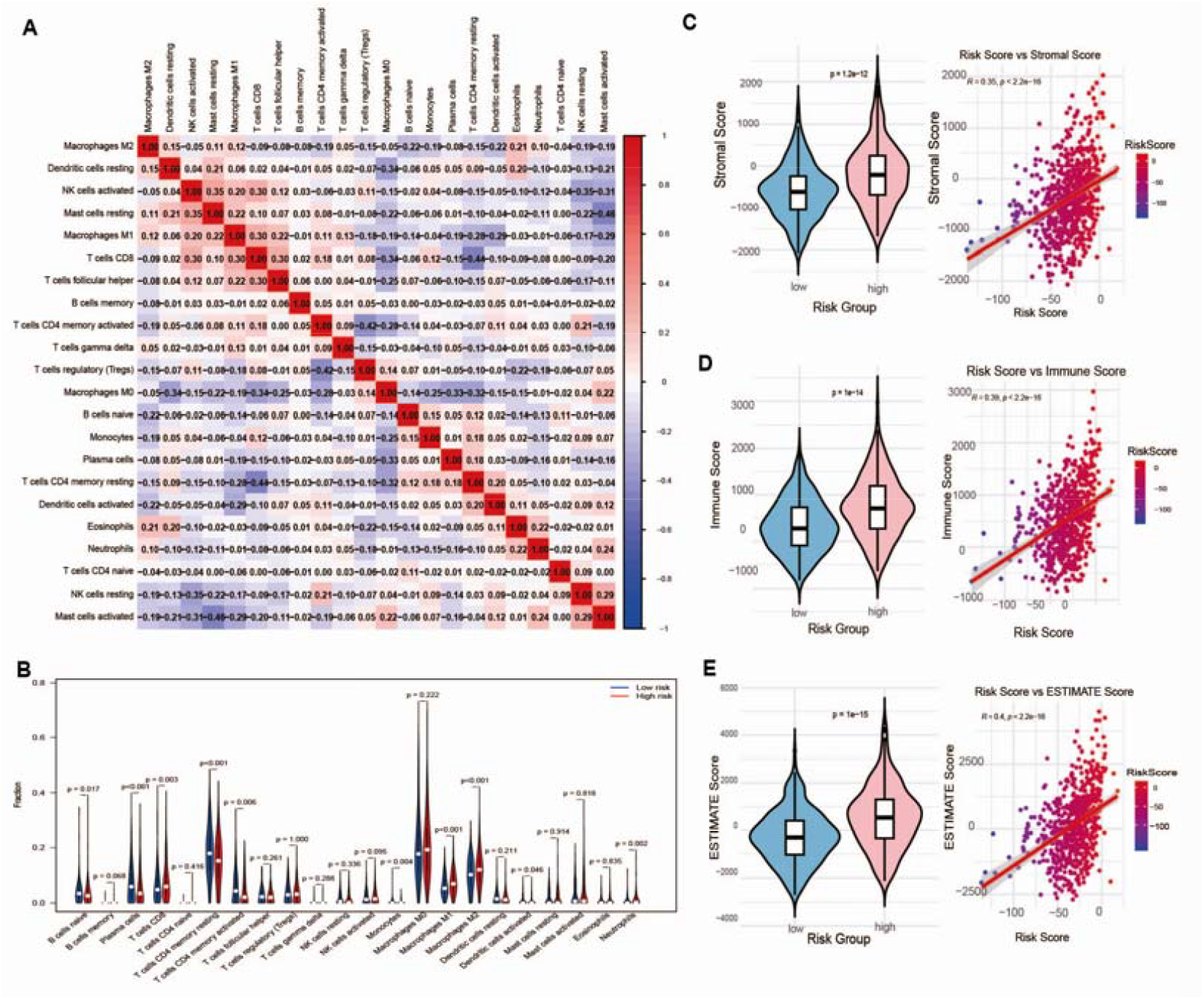
Immune cell infiltration profiles associated with the 8-gene signature. (A) Correlation matrix of 22 immune cell subsets via CIBERSORT. (B) Immune cell fraction comparison between high- and low-risk groups. (C–E) Violin plots and correlations between risk score and stromal, immune, and ESTIMATE scores.

Single-gene immune infiltration analysis revealed distinct correlations for key signature genes (Figure 6A). High AXIN2 expression was associated with increased anti-tumor immune cells, including naive B cells (*P* = 0.021), plasma cells (*P* = 0.003), and activated CD4^+^ memory T cells (*P* = 0.026) (Figure 6B), indicating a more activated anti-tumor immune microenvironment, consistent with its protective role. In contrast, high PSORS1C1 expression correlated with higher resting CD4^+^ memory T cells (*P* = 0.047) and trends toward reduced M1 macrophages (*P* = 0.011) and activated CD4^+^ memory T cells (*P* < 0.001) (Figure 6D), reflecting a pro-tumorigenic, immunosuppressive phenotype, consistent with its risk-factor role and poor prognosis association.

**Figure 6.**
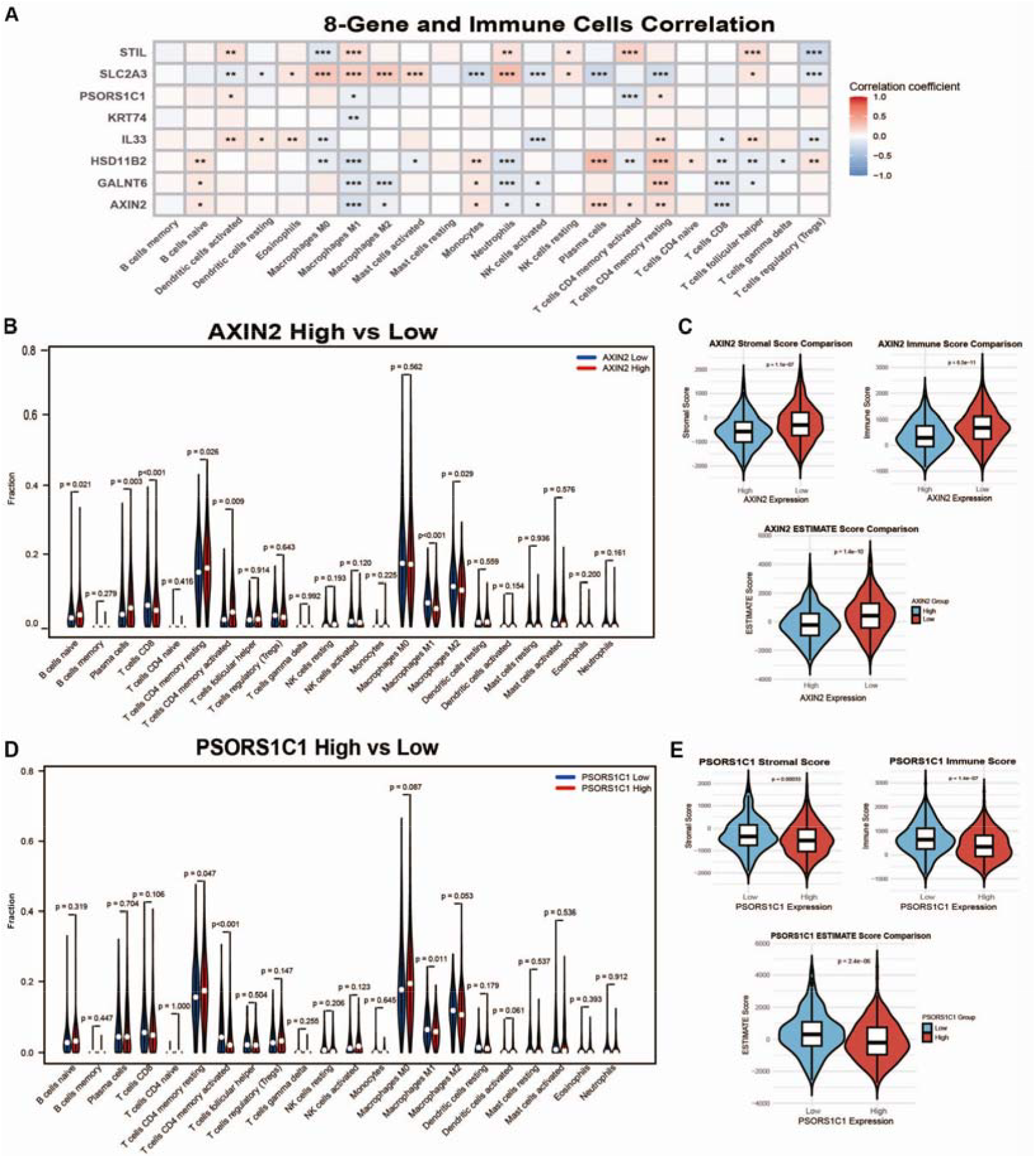
Immune correlations of key genes AXIN2 and PSORS1C1. (A) Heatmap of correlations between eight signature genes and 22 immune cell subsets. (B) Immune cell fractions in AXIN2 high/low groups. (C) Stromal, immune, and ESTIMATE scores in AXIN2 high/low groups. (D) Immune cell fractions in PSORS1C1 high/low groups. (E) Stromal, immune, and ESTIMATE scores in PSORS1C1 high/low groups.

### 3.6 ESTIMATE Immune Microenvironment Score Analysis

The ESTIMATE algorithm comprehensively evaluated immune and stromal characteristics. High-risk patients had significantly higher immune, stromal, and ESTIMATE scores (Figure 5C–E), indicating extensive stromal activation and complex immune infiltration. Correlation analysis confirmed positive associations between the risk score and all three microenvironment scores, showing the signature effectively reflects stromal and immune status. High AXIN2 expression correlated with a cleaner anti-tumor microenvironment (Figure 6C), while PSORS1C1 overexpression promoted stromal hyperplasia and immunosuppressive microenvironment formation (Figure 6E).

### 3.7 GSEA Revealed Immune-Resistant Pathway Activation

GSEA explored molecular mechanisms of bevacizumab resistance. Immune suppression-related pathways were most significantly enriched in the high-risk group (Figure 7A), with prominent activation of TNF-α/NF-κB, IL-6/JAK/STAT3, and immune checkpoint pathways, key mediators of inflammation, immune escape, and anti-angiogenic resistance (Figure 7D). Drug metabolism, DNA repair, and cell cycle pathways showed milder enrichment. Single-gene GSEA confirmed opposing immune regulation by AXIN2 and PSORS1C1: AXIN2 suppressed inflammatory and immune checkpoint pathways (Figure 7B,E), while PSORS1C1 activated multiple immunosuppressive cascades (Figure 7C,F). PSORS1C1 also regulated cell cycle and DNA damage repair, indicating multi-dimensional control of proliferation and resistance. Immune regulation was the core mechanism of the signature in bevacizumab resistance.

**Figure 7.**
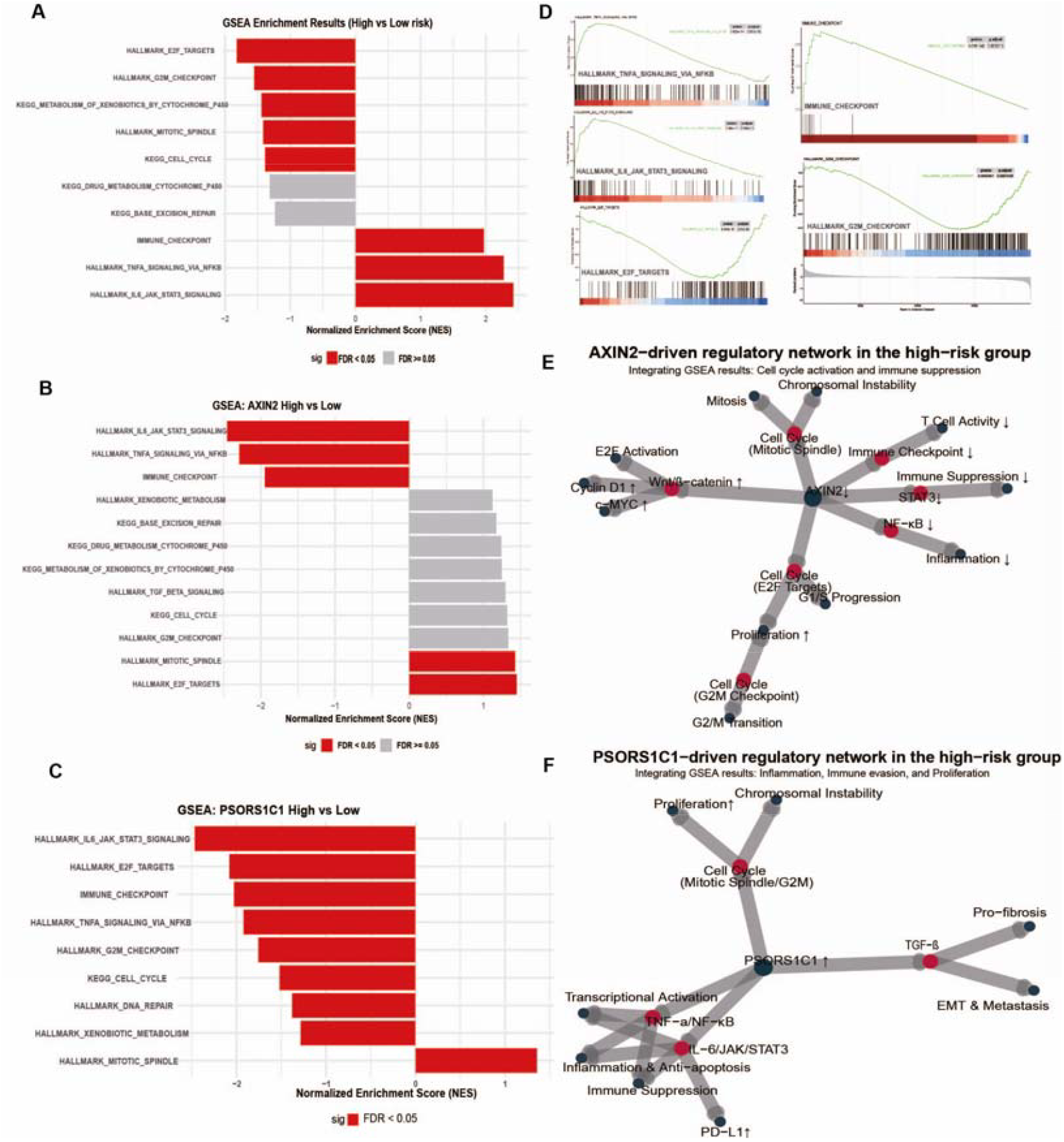
GSEA and functional regulatory network analysis. (A–C) GSEA results for high-risk, AXIN2 high, and PSORS1C1 high groups. (D) Enrichment plots of representative pathways (TNF-α/NF-κB, IL-6/JAK/STAT3, immune checkpoint, E2F targets) in high-risk patients. (E) AXIN2-driven regulatory network integrating cell cycle, immune suppression, and inflammation pathways. (F) PSORS1C1-driven regulatory network highlighting inflammation, immune evasion, proliferation, and metastasis pathways.

## 4 Discussion

Bevacizumab resistance remains a major challenge in metastatic CRC [11,12]. The immune microenvironment critically influences therapeutic resistance [13,14]. Here, an 8-gene bevacizumab resistance-related prognostic signature was developed and validated, with systematic exploration of immune infiltration and molecular pathways.

First, the signature was constructed from overlapping DEGs of bevacizumab-resistant transcriptomes and TCGA tumor profiles, ensuring linkage to both malignancy and drug resistance. It showed stable prognostic performance in TCGA and GSE39582 cohorts, with the risk score confirmed as an independent prognostic factor. Compared with prior models [5–7], this signature has higher translational potential due to direct linkage to bevacizumab resistance.

Second, high-risk CRC was characterized by immunosuppression linked to bevacizumab resistance. High-risk patients had increased M2 macrophages, which secrete VEGF, IL-10, and TGF-β to drive pathological angiogenesis, impair vascular normalization, and counteract bevacizumab efficacy [15,16]. Elevated M1/M0 macrophages reflect dysregulated inflammation: M1-derived IL-6/TNF-α promotes angiogenesis and immune suppression, while M0 expansion favors M2 polarization [15,16]. Despite higher CD8^+^ T cells, reduced activated dendritic cells and memory CD4^+^ T cells indicate impaired cytotoxic immunity [17,18]. Increased neutrophils amplify resistance via ROS, elastase, and pro-angiogenic factors, synergizing with M2 macrophages [19,20]. Reduced monocytes, activated dendritic cells, and helper T cells confirm blunted adaptive immunity [18]. Collectively, high-risk CRC exhibits M2 polarization, myeloid inflammation, and defective cytotoxic immunity, supporting combined anti-angiogenic and immunomodulatory therapy.

Third, AXIN2 and PSORS1C1 were identified as core independent prognostic factors with opposing roles in shaping the TIME and bevacizumab resistance. AXIN2, a negative Wnt/β-catenin regulator [21,22], correlated with increased anti-tumor immune cells and a “hot” microenvironment, restraining excessive inflammation and immune suppression. PSORS1C1 correlated with immunosuppressive cell accumulation, promoting a pro-tumorigenic phenotype [23,24]. These divergent profiles align with their prognostic roles: AXIN2 as a protective factor, PSORS1C1 as a risk factor. Their imbalance drives immunosuppressive microenvironment transition, contributing to bevacizumab resistance and poor outcomes.

Finally, GSEA highlighted enrichment of TNF-α/NF-κB, IL-6/JAK/STAT3, and immune checkpoint pathways in the high-risk group, forming a positive feedback loop for immune escape and resistance [25–29]. AXIN2 and PSORS1C1 synergistically regulated immune suppression-related pathways and modulated cell cycle, DNA repair, and drug metabolism, forming a multi-pathway regulatory network for bevacizumab resistance [30,31]. Immune suppression was the core resistance hub, with other pathways playing auxiliary roles.

This study has limitations. It is retrospective, based on public databases, requiring prospective validation. Further in vitro and in vivo experiments are needed to verify AXIN2 and PSORS1C1 functions in bevacizumab resistance.

## 5 Conclusion

An 8-gene bevacizumab resistance-related prognostic signature was constructed and validated, effectively predicting CRC prognosis and reflecting the immunosuppressive microenvironment linked to bevacizumab resistance. AXIN2 and PSORS1C1 are core independent prognostic factors and key regulators of immune-mediated resistance. This signature provides a promising tool for prognostic stratification, therapeutic decision-making, and mechanistic exploration in CRC.

## Acknowledgments

This study was supported by the Clinical Research Special Project of Shanghai Municipal Health Commission (Grant No. 20214Y0141). The authors sincerely thank all participants and researchers who contributed to the public databases (TCGA and GEO) for providing open access data. We also gratefully acknowledge the technical support and academic guidance from colleagues in our research group during this study.

## Author Contributions

Niu ZC designed the study, supervised the research, and revised the manuscript. Qiu DZ performed data analysis, drafted the original manuscript, and prepared figures. Xu PP provided clinical resources, interpreted the results, and critically revised the manuscript. All authors read and approved the final version.

---

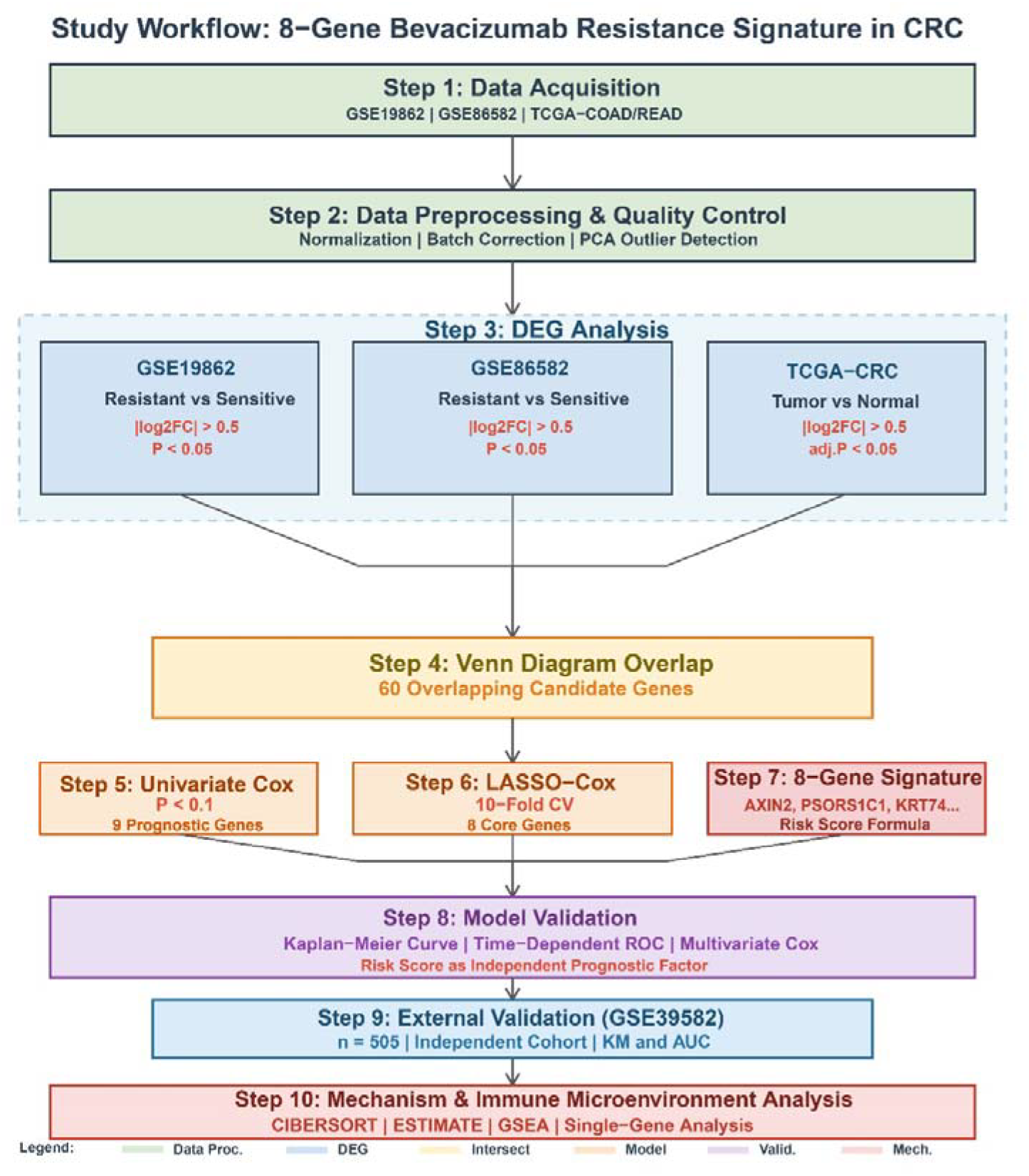

**Flowchart** Study workflow diagram. The step-by-step design of this study, including data acquisition, DEG analysis, signature construction, validation, and functional exploration in CRC patients.

